# Homeostatic responses to hypoxia by the carotid body and adrenal medulla are based on mutual antagonism between HIF-1α and HIF-2α

**DOI:** 10.1101/2022.07.11.499380

**Authors:** Guoxiang Yuan, Ying-Jie Peng, Vaddi Damodar Reddy, Vladislav Makarenko, Jayasri Nanduri, Shakil A. Khan, Joseph A. Garcia, Ganesh K. Kumar, Gregg L. Semenza, Nanduri R. Prabhakar

**Author notes:** Contributed equally to this work (Guoxiang Yuan, Ying-Jie Peng, Vaddi Damodar Reddy, Vladislav Makarenko). Author contributions: N.R.P. designed research; G.Y., Y.-J.P., V.D.R., V.M., J.N., S.A.K., and G.K.K. performed research; J.A.G. and G.L.S. contributed reagents/analytical tools; G.Y., Y.-J.P., V.D.R., V.M., J.N., S.A.K., and G.K.K. analyzed the data; N.R.P., G.K.K.,and G.L.S. wrote the paper.

## Abstract

Respiration and blood pressure (BP) are regulated to maintain optimal delivery of O_2_ to every cell in the body. Arterial hypoxemia is sensed by the carotid body (CB), which initiates sympathetic reflex arcs to the diaphragm to increase ventilation, and to the adrenal medulla (AM) to increase catecholamine secretion and thereby increase BP. However, the underlying molecular mechanisms have not been fully delineated. Here, we report that the relative activities of hypoxia-inducible factor-1 (HIF-1) and HIF-2 determine the set point for the CB and AM, with respect to their maintenance of BP and respiration. In *Hif2a^+/-^* mice, which are heterozygous for a knockout allele at the locus encoding HIF-2α, expression of HIF-1α and NADPH oxidase 2 was increased in the CB and AM, resulting in an oxidized intracellular redox state with augmented sensitivity to hypoxia, increased BP, and respiratory abnormalities, which were all normalized by treatment with a HIF-1α inhibitor or a superoxide anion scavenger. By contrast, in *Hif1a^+/-^* mice, which are heterozygous for a knockout allele at the locus encoding HIF-1α, the expression of HIF-2α and superoxide dismutase 2 was increased in the CB and AM, resulting in a reduced intracellular redox state with impaired CB and ventilatory responses to chronic hypoxia, which were normalized by treatment with a HIF-2α inhibitor. None of the abnormalities that were observed in *Hif1a^+/-^* or *Hif2a^+/-^* mice were observed in *Hif1a^+/-^*; *Hif2a^+/-^* double-heterozygous mice. Our results demonstrate that redox balance in the CB and AM, which is determined by mutual antagonism between HIF-α isoforms, establishes the set point for responses of the CB and AM to hypoxia, and is required for the maintenance of normal BP and respiration.

## Introduction

The respiratory and cardiovascular systems of vertebrates are designed to insure optimal O_2_ delivery to every cell at every moment of the organism’s lifetime. O_2_ is essential for survival because it is required to generate sufficient ATP to maintain the structure and function of large, complex organisms (1). During the oxidation of acetyl CoA derived from pyruvate or fatty acids in the tricarboxylic acid cycle, the energy trapped in carbon-carbon bonds is used to reduce NAD^+^ to NADH. Electrons from NADH flow through the electron transport chain (before reacting with O_2_ in complex IV to form water), establishing a proton gradient that is used to synthesize ATP in complex V. Under conditions of decreased O_2_ availability (i.e. hypoxia), the conversion of pyruvate to acetyl CoA is inhibited, thereby decreasing NADH and ATP generation. Thus, cellular O_2_ availability is critical for energy and redox homeostasis. O_2_ delivery in turn depends on proper regulation of breathing and blood pressure (BP).

The carotid body (CB) is the sensory organ that stimulates breathing in response to hypoxemia, thereby ensuring adequate systemic O_2_ availability (2). Hypoxia also directly stimulates chromaffin cells in the adrenal medulla (AM) to increase catecholamine secretion, in both neonates (3–6) and adults (7), leading to increased BP, thereby facilitating tissue O_2_ delivery. Despite decades of study, the molecular mechanisms that establish the set point for CB and AM function under normoxic conditions, and result in augmented activity under hypoxic conditions, have not been delineated.

The discovery of hypoxia-inducible factor 1 (HIF-1), and subsequent identification of HIF-2, provided a paradigm shift in our understanding of oxygen homeostasis (8). In HIF-1 and HIF-2, an O_2_-labile HIF-1α or HIF-2α subunit, respectively, dimerizes with HIF-1β. HIF-1α is ubiquitously expressed in all animals, whereas HIF-2α is a product of vertebrate evolution (9). Homozygosity for a knockout allele at the locus encoding HIF-1α or HIF-2α leads to embryonic lethality, whereas mice that are heterozygous for a knockout allele develop normally (10, 11).

HIF-1α and HIF-2α are expressed in the CB (12, 13), but the functional consequences of their expression have not been delineated. Although both HIF-1α and HIF-2α are both positive regulators of angiogenesis and erythropoiesis (14), CB responses to hypoxia are impaired in *Hif1α^+/-^* mice (15), whereas they are augmented in *Hif2α^+/-^* mice (16). The mechanisms underlying this striking contrast in CB responses to hypoxia in *Hif1α^+/-^* and *Hif2α^+/-^* mice are not known. Consistent with the contrasting phenotypes of the heterozygous knockout mice, HIF-1 and HIF-2 regulate genes with opposing functions in the CB. HIF-1 is a positive regulator of Nox2, which encodes the pro-oxidant enzyme NADPH oxidase 2 (17). By contrast, HIF-2 is a positive regulator of *Sod2*, which encodes the antioxidant enzyme superoxide dismutase 2 (11, 18). Based on these findings, we tested the hypothesis that the relative expression of HIF-1α and HIF-2α regulates redox state in the CB and AM, and by doing so, determines the set point for changes in breathing and BP in response to changes in arterial O_2_ levels.

## Results

### HIF-1α levels are increased in the CB and AM of *Hif2α^+/−^* mice

Tissue sections from *Hif2α^+/−^* and wild type (WT) littermates were analyzed by immunofluorescence, utilizing antibodies against HIF-1α and chromogranin A (CGA), which is expressed specifically in the O_2_-sensing glomus cells of the CB (19). Compared to WT mice, HIF-1α protein levels were markedly increased in the CB of *Hif2α^+/−^* mice (Figure 1A). Immunoblot (IB) assays revealed that HIF-1α expression was also increased in the AM from *Hif2α*^+/−^ mice (Figure 1B). By contrast, HIF-1α mRNA expression was not increased in the CB or AM of *Hif2α*^+/−^ mice (Figure S1A-B).

**Figure 1.**
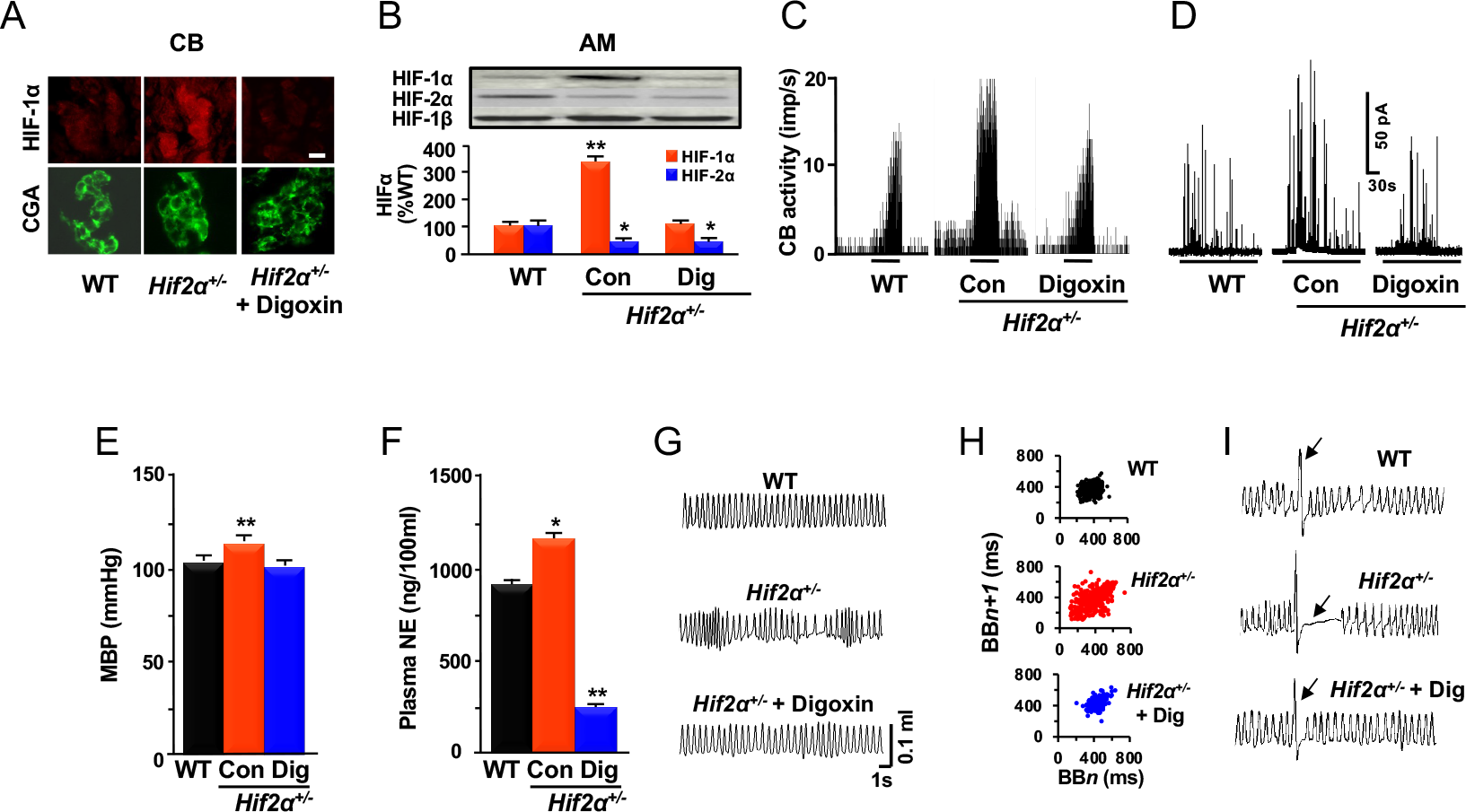
Analysis of *Hif2α*^+/−^ mice. Wild type mice (WT), control (Con) *Hif2α^+/−^* mice, and *Hif2α^+/−^* mice treated with digoxin (Dig) were studied. (**A**) Immunofluorescence assay for HIF-1α and chromogranin A (CGA) in the carotid body (CB). Scale bar = 20 μm. (**B**) Immunoblot (IB) assays (*top*) and densitometric analysis (*bottom*) of HIF-1α (*red*) and HIF-2α (*blue*) protein levels in the adrenal medulla (AM) relative to untreated WT mice are shown. (**C**) CB neural activity in response to hypoxia (*P_O_*_2_ = 40 mm Hg for 3 minutes as indicated by bar under tracing) is shown. Activity is presented as impulses per second (imp/s). (**D**) Catecholamine secretion from AM chromaffin cells in response to hypoxia (*PO2* = 40 mm Hg as indicated by bar under tracing) was measured by carbon fiber amperometry. (**E** and **F**) Mean blood pressure (MBP; **E**) and plasma norepinephrine (NE) levels (**F**) were determined. (**G-I**) Breathing pattern (**G**), Poincarè plots of breath-to-breath (BB) interval in milliseconds (**H)**, and post-sigh apneas (arrows; **I**) were analyzed. Data are presented as mean ± SEM, *n* = 6-8 mice per group; \**p* < 0.05, \*\**p* < 0.01.

Digoxin inhibits the accumulation of HIF-1α protein (20) and after daily intraperitoneal (IP) injection of digoxin (1 mg/kg) for three days, HIF-1α levels were no longer increased in the CB (Figure 1A) or AM (Figure 1B) of *Hif2α^+/−^* mice. CB and AM responses to hypoxia were evaluated in *Hif2α^+/−^* mice treated with digoxin or vehicle, as described previously (16, 21). Compared to WT mice, *Hif2α^+/−^* mice that were treated with vehicle were found to have increased CB neural activity (Figures 1C and S1C) and enhanced catecholamine secretion from AM chromaffin cells in response to hypoxia (Figures 1D and S1D). Mean BP (Figure 1E) and plasma norepinephrine (NE) levels (Figure 1F) were increased in *Hif2α^+/−^* mice. The animals also showed signs of breathing instability, including periods of hypoventilation followed by hyperventilation and post-sigh apneas, which were defined as cessation of breathing for a duration greater than three breaths (Figures 1G-I and S1E-G). Bilateral transection of the carotid sinus nerve eliminated the respiratory abnormalities in *Hif2α^+/−^* mice (Figure S1H-L), indicating the presence of augmented CB neural activity under normoxic conditions. The augmented hypoxic responses of the CB and AM, hypertension, and ventilatory abnormalities were also eliminated in digoxin-treated *Hif2α^+/−^* mice (Figures 1C-I and S1C-L), in which HIF- 1α levels were decreased to levels similar to untreated WT littermates (Figure 1A-B).

### HIF-2α levels are increased in the CB and AM of *Hif1α^+/−^* mice

Compared to WT mice, HIF-2α expression was increased in CB (Figure 2A) and AM (Figure 2B) of *Hif1α^+/−^* mice. Increased HIF-2α expression in the CB and AM was not due to increased mRNA levels (Figure S2A-B). After demonstrating that treatment of PC12 pheochromocytoma cells (derived from a tumor of AM chromaffin cells) with 2-methoxyestradiol decreased HIF- 2α expression (Figure S2C), we administered 2-methoxyestradiol to *Hif1α^+/−^* mice (200 mg/kg/day IP for 3 days), which decreased HIF-2α protein levels in the CB (Figure 2A) and AM (Figure 2B) to WT levels. The increased HIF-2α expression in *Hif1α^+/−^* mice was associated with impaired CB and AM chromaffin cell responses to hypoxia, which were normalized following treatment with 2-methoxyestradiol (Figures 2C-D and S2D-E).

**Figure 2.**
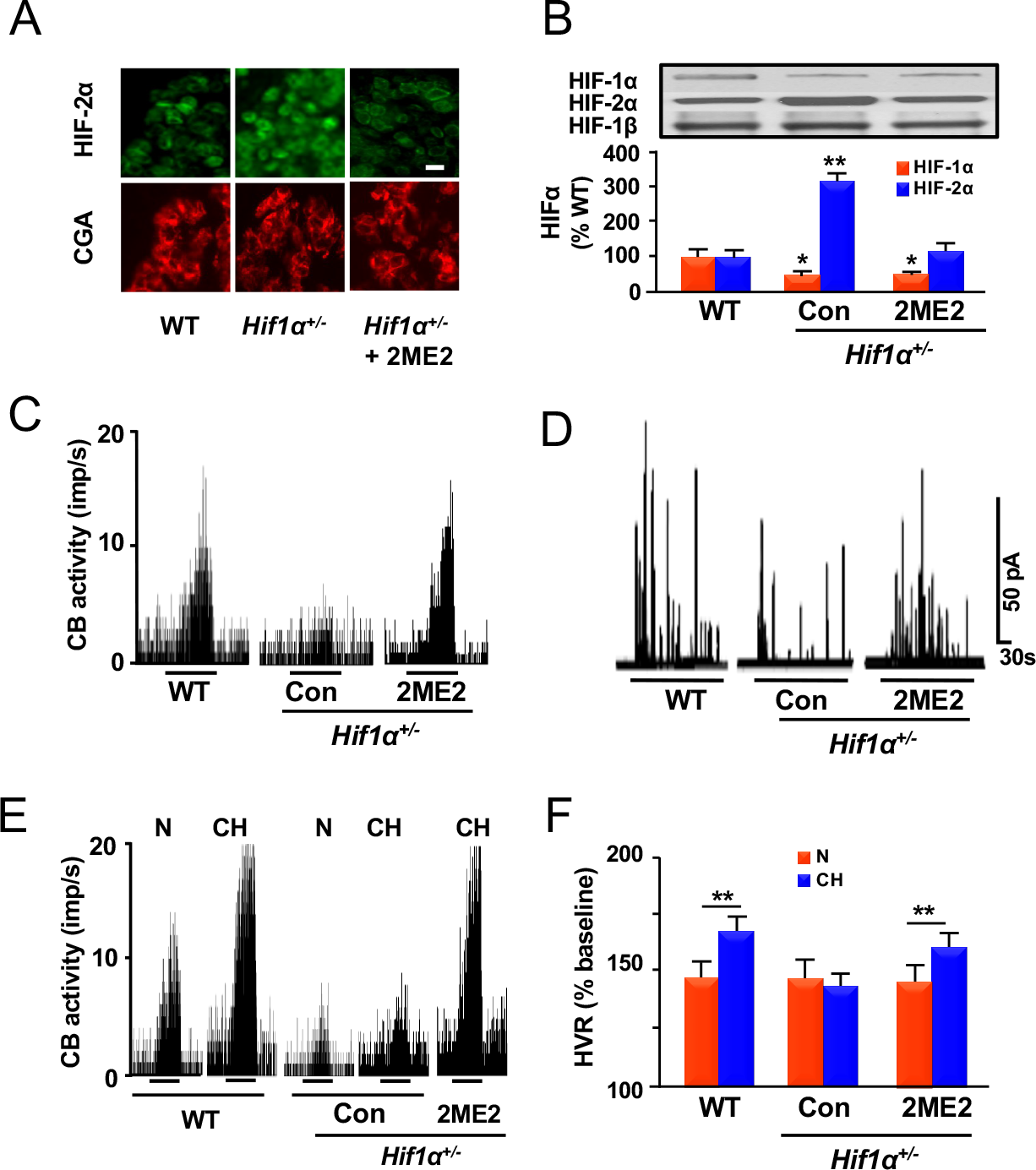
Analysis of *Hif1α*^+/−^ mice. WT type mice, control untreated *Hif1α^+/−^* mice (Con), and *Hif1α^+/−^* mice treated with 2-methoxyestradiol were studied. (**A**) HIF-2α and CGA expression in the CB was analyzed by immunofluorescence. Scale bar = 20 μm. (**B**) IB assays (*top*) and densitometric analysis (*bottom*) of HIF-1α and HIF-2α protein. expression in the AM relative to untreated WT mice are shown. (**C**) CB responses to hypoxia (*P_O_*_2_ = 40 mmHg for 3 minutes as indicated by black bar). (**D**) Catecholamine secretion from AM chromaffin cells in response to hypoxia (black bar). (**E**) Responses to hypoxia were analyzed in CBs from mice maintained under normoxic conditions (N) or exposed to chronic hypoxia (CH; 72 hours at 0.4 atmospheres). (**F**) The hypoxic ventilatory response (HVR) after CH was analyzed as ratio of minute ventilation (V_E_) to O_2_ consumption (V_O2_) and presented as percent of baseline (prior to CH). Data in graphs are presented as mean ± SEM, *n* = 7-8 mice; \**p* < 0.05, \*\**p* < 0.01.

Mice subjected to chronic hypoxia manifest augmented CB responses to a subsequent acute decrease in *p*O_2_, a phenomenon known as ventilatory acclimatization (19). WT mice exposed to chronic hypobaric hypoxia (0.4 atmospheres for 72 hours) showed an increased ventilatory response to a subsequent acute hypoxic exposure; by contrast, ventilatory acclimatization was lost in *Hif1α^+/−^* mice (15). However, treatment with 2-methoxyestradiol to normalize HIF-2α levels restored the CB (Figures 2E and S2F) and ventilatory (Figure 2F) response in *Hif1α^+/−^* mice following exposure to chronic hypoxia.

### Inhibition of HIF-α subunits alters CB and AM function in WT mice

Compared to vehicle treatment, digoxin treatment of WT mice led to decreased HIF-1α and increased HIF-2α expression in the AM (Figure S3A), decreased hypoxia-induced CB activity (Figure S3B-C), and decreased NE levels (Figure S3D), which were all features similar to the phenotype of *Hif1α^+/−^* mice (Figure 2). Likewise, similar to *Hif1α^+/−^* mice, digoxin-treated WT mice had no BP or breathing abnormalities (Figure S3E-H).

Treatment of WT mice with 2-methoxyestradiol, which inhibits HIF-2α expression (Figure S2C), increased hypoxia-induced CB neural activity (Figure S4A-B), irregular breathing (Figure S4C-E), increased post-sigh apneas (Figure S4F-G), an increase in mean BP of 8 ± 1.2 mm Hg (Figure S4H), and greatly increased NE levels (Figure S4I), which were similar to the phenotype of *Hif2α^+/−^* mice (Figure 1).

### Analysis of *Hif1α^+/-^*; *Hif2α^+/-^* double heterozygous (DH) mice

We hypothesized that the phenotypes of *Hif1α^+/-^* and *Hif2α^+/-^* mice were not due to decreased absolute levels of HIF- 1α and HIF-2α, respectively, but rather due to changes in the levels of HIF-1α and HIF-2α relative to each other. To test this hypothesis, we generated *Hif1α^+/-^*;*Hif2α^+/-^* DH mice. Both HIF-1α and HIF-2α protein levels were significantly but equally reduced in the AM from *Hif1α^+/-^*; *Hif2α^+/-^* DH mice (Figure 3A). Despite the deficiency of both HIF-1α and HIF-2α, the CB and AM responses to acute hypoxia (Figures 3B-C and S5A-B), BP (Figure 3D), NE levels (Figure 3E), breathing pattern (Figure 3F and S5C-E), as well as the CB and ventilatory responses to chronic hypoxia (Figures 3G-H and S5F) in DH mice were all indistinguishable from WT littermates.

**Figure 3.**
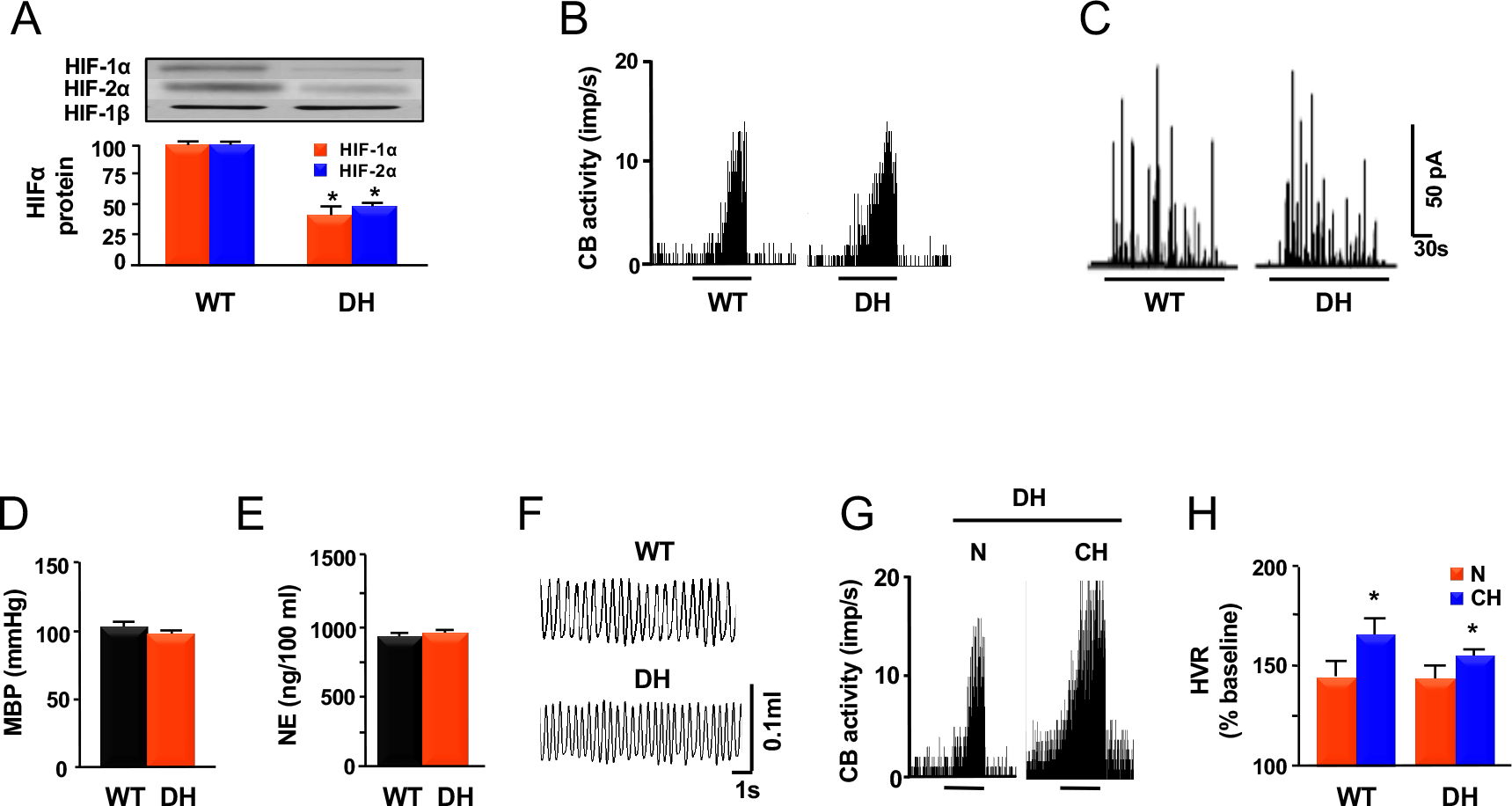
Analysis of double-heterozygous (DH) mice. WT (*Hif1α^+/+^;Hif2α^+/+^*) and DH (*Hif1α^+/-^;Hif2α^+/-^*) mice were studied. (**A**) IB assays (*top*) and densitometric analysis (*bottom;* mean ± SEM, *n* = 6 mice each) of HIF-1α and HIF-2α expression in the AM from DH and WT mice are shown. (*B*) CB responses to hypoxia are shown. (**C**) AM chromaffin cell catecholamine secretory response to hypoxia (black bar) was analyzed. (**D** and **E**) MBP (**D**) and plasma NE levels (**E**) are shown. (**F**) Breathing patterns of WT and DH mice. (**G**) Response to hypoxia in CBs from mice maintained under normoxic conditions (N) and mice exposed to chronic hypoxia (CH). (**H**) Hypoxic ventilatory response (HVR) after 72 hours of CH was analyzed as ratio of V_E_ to V_O2_ and percent of baseline is presented as mean ± SEM from 6 mice. **\****p* < 0.05 (graph in **A** and **H**).

### Cellular redox state is altered in the AM of *Hif2α^+/−^* and *Hif1α^+/−^* mice

We previously reported that *Hif2α^+/−^* mice exhibit oxidative stress and that antioxidant treatment corrected their cardio-respiratory abnormalities (16). Oxidative stress in *Hif2α^+/−^* mice was attributed to decreased expression of genes encoding antioxidant enzymes, including *Sod2* (11, 16, 18). By contrast, HIF-1α is a positive regulator of the *Nox2* gene (17). We hypothesized that HIF-1α-dependent Nox2 expression promotes oxidative stress in *Hif2α^+/-^* mice. Nox2 and Sod2 mRNA levels and enzyme activities were determined in the AM. Because aconitase is inactivated by oxidation (22), we determined aconitase catalytic activity as a measure of oxidative stress. These analyses were performed using AM because CB tissue (∼25 μg per mouse) is insufficient for biochemical assays.

Nox2 mRNA and enzyme activity were increased in AM from untreated *Hif2α^+/-^* mice but not in digoxin-treated *Hif2α^+/-^* mice (Figure 4A). By contrast, Sod2 mRNA expression and enzyme activity were decreased in AM from both control and digoxin-treated *Hif2α^+/-^* mice (Figure 4B). Aconitase activity was decreased in cytosolic and mitochondrial fractions from AM of control *Hif2α^+/-^* mice, and aconitase activity was normalized by digoxin treatment (Figure 4C).

**Figure 4.**
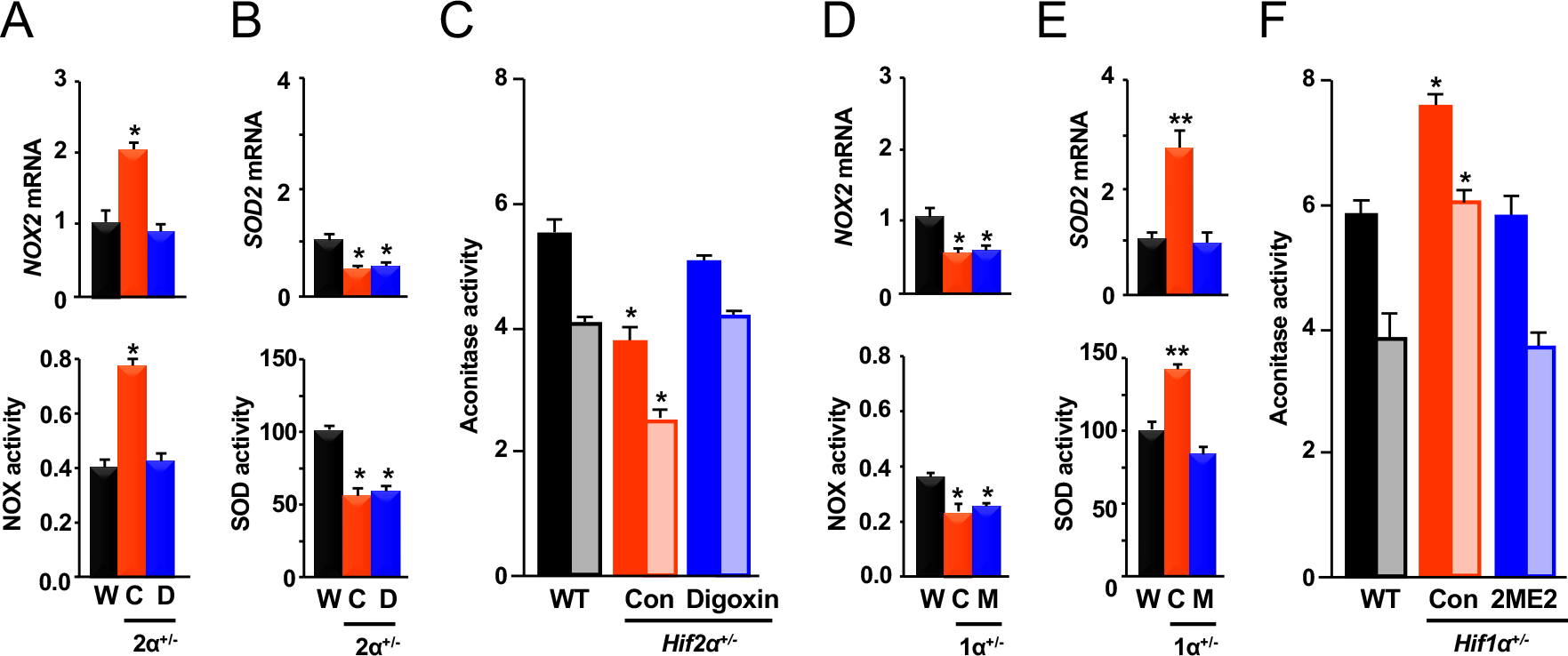
Analysis of intracellular redox state in the AM of *Hif2α^+/-^* and *Hif1α^+/-^* mice. NADPH oxidase 2 (Nox2) and superoxide dismutase 2 (Sod2) mRNA and enzyme activity, and cytosolic (*dark bars*) and mitochondrial (*shaded bars*) aconitase activities were determined in AM from wild type mice (W or WT), control (C or Con) *Hif1α^+/−^* or *Hif2α^+/−^* mice, *Hif2α^+/−^* mice treated with digoxin (D), and *Hif-1α^+/−^* mice treated with 2- methoxyestradiol (M). (**A-C**) Data from WT, *Hif2α^+/−^* and *Hif2α^+/−^* + digoxin groups. (**D-F**) Data from WT, *Hif1α^+/−^* and *Hif1α^+/−^* + 2-methoxyestradiol groups. Data in graphs are presented as mean ± SEM, *n* = 6-8 mice per group; \**p* < 0.05, \*\**p* < 0.01.

Analysis of AM from *Hif1α^+/-^* mice revealed decreased Nox2 mRNA and catalytic activity (Figure 4D), and increased Sod2 mRNA and catalytic activity (Figure 4E). Compared to WT littermates, aconitase catalytic activity was increased in cytosolic and mitochondrial fractions of AM from *Hif1α^+/-^* mice (Figure 4F). These findings indicate that partial HIF-1α deficiency leads to a reduced intracellular redox state in the AM. Treatment of *Hif1α^+/-^* mice with 2-methoxyestradiol, which normalized HIF-2α expression (Figure 2B), also normalized Sod2 expression and activity (Figure 4E) and aconitase activity in cytosolic and mitochondrial fractions of AM (Figure 4F). However, 2-methoxyestradiol treatment had no effect on Nox2 mRNA expression or catalytic activity in AM of *Hif1α^+/-^* mice (Figure 4D). Treatment with 2-methoxyestradiol increased Nox2 and decreased Sod2 protein expression in AM from WT mice (Figure S4K). In contrast to the altered redox states demonstrated in AM from *Hif1α^+/-^* and *Hif2α^+/-^* mice, analysis of AM from DH mice revealed that Nox2 and Sod2 mRNA levels and catalytic activities, as well as cytosolic and mitochondrial aconitase activities, were not significantly different from WT littermates (Figure S5G-I).

### mTOR and prolyl hydroxylase activity is altered in HIF-2α-deficient PC12 cells

We next investigated the molecular mechanisms by which partial HIF-2α deficiency led to increased HIF-1α expression. These studies utilized PC12 cells because they involved silencing or forced expression of HIF-2α, which are technically challenging in primary cultures of CB glomus cells or AM chromaffin cells. PC12 cells were chosen because they are derived from the AM and respond to hypoxia with increased catecholamine secretion (23) similar to AM chromaffin and CB glomus cells (19). Treatment of PC12 cells with a small interfering RNA (siRNA) targeted against HIF-2α (siHIF-2α) led to decreased HIF- 2α and Sod2 expression, increased HIF-1α and Nox2 expression, and decreased aconitase activity as compared to untreated control cells or cells treated with a scrambled control siRNA (Figure 5A-B), which was similar to the phenotype observed in AM from *Hif-2*α*^+/-^* mice (Figure 4A-C). Forced expression of Sod2 in siHIF-2α-treated cells relieved the oxidative stress and prevented the increase in HIF-1α expression (Figure 5B), suggesting that oxidative stress triggers increased HIF-1α expression in HIF-2α-deficient PC12 cells. We next investigated how oxidative stress increases HIF-1α protein levels in HIF-2α-deficient PC12 cells. We have previously reported that oxidative stress induced by intermittent hypoxia increases HIF-1α protein synthesis and that this effect was mediated by activation of mammalian target of rapamycin (mTOR), a kinase that stimulates translation of HIF-1α mRNA (24). Compared to cells treated with scrambled siRNA, phosphorylated (active) mTOR levels were increased in siHIF-2α−treated cells, whereas the levels of phospho-mTOR and HIF-1α were normalized when oxidative stress was relieved either by treatment with MnTMPyP, a membrane permeable antioxidant, or by forced Sod2 expression (Figure 5*C*). Treatment of siHIF-2α−treated cells with rapamycin, a selective mTOR inhibitor, eliminated phospho-mTOR and decreased HIF-1α expression by approximately 60% (Figure 5C), indicating mTOR promotes increased HIF-1α expression in HIF-2α-deficient cells.

**Figure 5.**
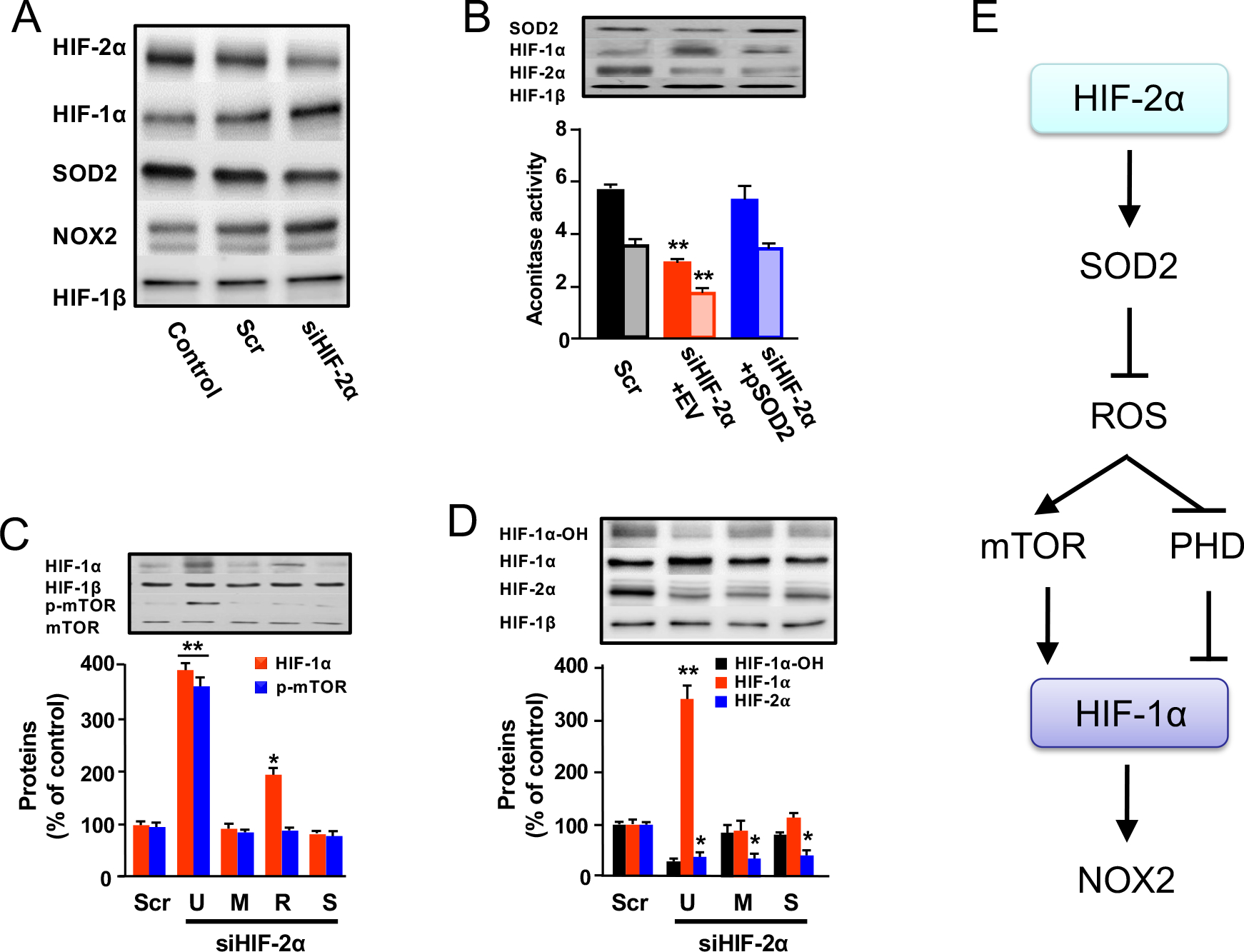
Regulation of HIF-1α expression by HIF-2α in PC12 cells. (**A**) Levels of the indicated proteins were determined in lysates of non-transfected control cells and in cells transfected with scrambled siRNA (Scr) or HIF-2α siRNA (siHIF-2α). (**B**) Cells were transfected with Scr or siHIF-2α and either empty vector (EV) or Sod2 expression vector (pSod2), followed by IB assays (*top*) and measurement of aconitase activity (*bottom*) in cytosolic (*solid bars*) and mitochondrial (*shaded bars*) fractions. (**C** and **D**) IB assays were performed using lysates of cells that were transfected with Scr or siHIF-2α and left untreated (U); treated with MnTMPyP (M; 50 μM) or rapamycin (R; 100 nM); or transfected with Sod2 expression vector (S). IB assays (*top*) and densitometric analysis (*bottom*) are shown. Data in graphs are presented as mean ± SEM, *n* = 4-5 independent experiments; \**p* < 0.05, \*\**p* < 0.01. (**E**) The mechanism by which HIF-2α negatively regulates HIF-1α and Nox2 expression is shown. Arrow, stimulatory effect; blocked arrow, inhibitory effect.

O_2_-dependent hydroxylation by prolyl hydroxylases triggers HIF-1α degradation, whereas oxidation of the Fe^2+^ ion in the catalytic center of these enzymes inactivates them (25). Levels of hydroxylated HIF-1α were decreased and total HIF-1α expression was increased in cells treated with siHIF-2α, whereas MnTMPyP treatment or forced Sod2 expression counteracted the effects of siHIF-2α (Figure 5D). Thus, both oxidant-dependent inhibition of prolyl hydroxylation and oxidant-dependent activation of mTOR contribute to increased HIF-1α expression in HIF-2α-deficient PC12 cells (Figure 5E).

### Altered Ca^2+^ levels and calpain activity in HIF-1α−deficient cells

We next investigated the molecular mechanisms that mediate increased HIF-2α expression in HIF-1α deficient cells. Treatment of PC12 cells with siHIF-1α led to decreased HIF-1α and Nox2 expression, increased HIF-2α and Sod2 expression, and a reduced intracellular redox state as evidenced by increased aconitase catalytic activity (Figure 6A-B). Forced expression of Nox2 in siHIF-1α−treated cells normalized aconitase catalytic activity and HIF-2α expression (Figure 6B).

**Figure 6.**
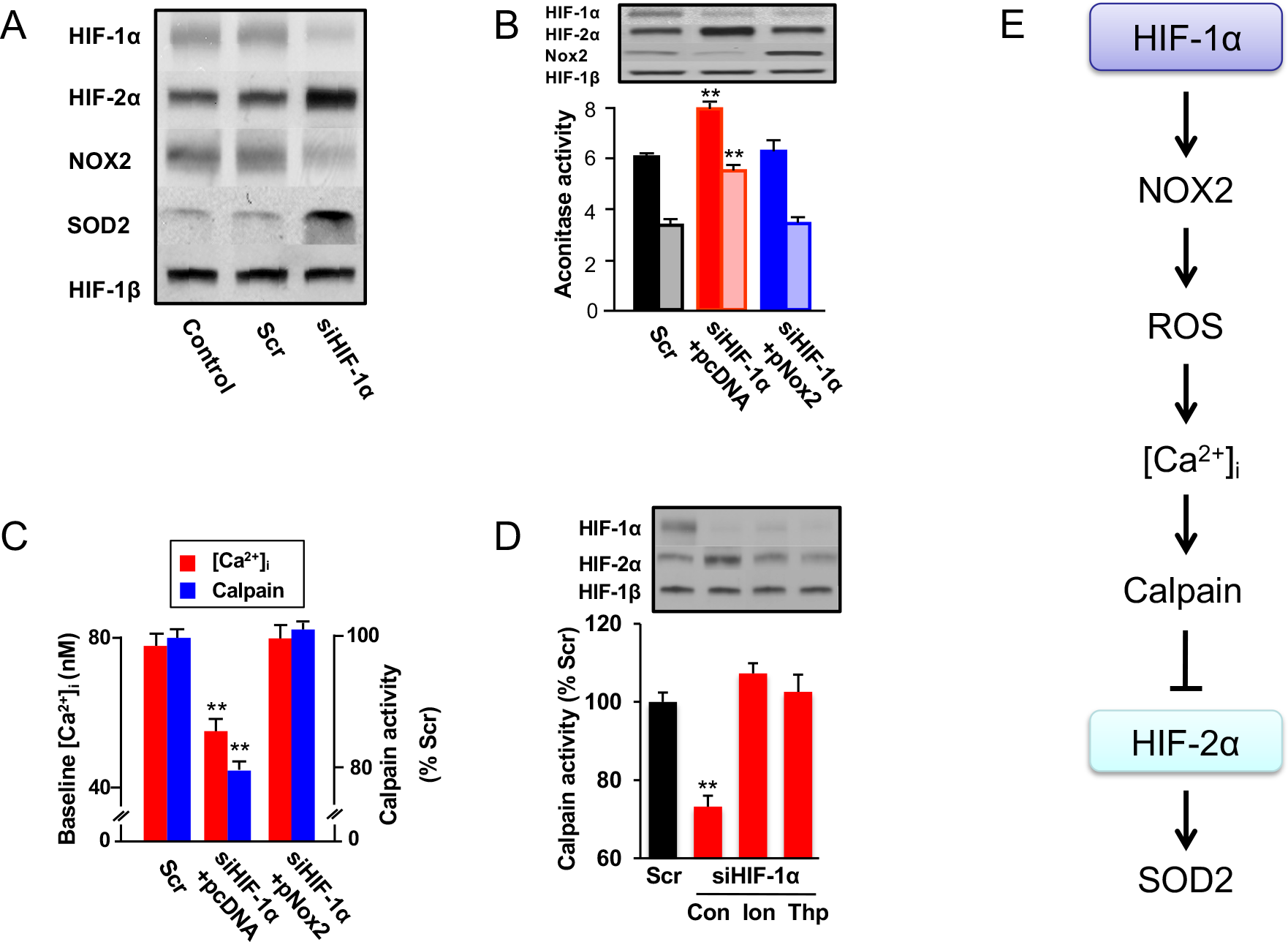
Regulation of HIF-2α expression by HIF-1α in PC12 cells. (**A**) Levels of the indicated proteins were determined in lysates of control cells and cells transfected with scrambled siRNA (Scr) or HIF-1α siRNA (siHIF-1α). (**B**) Cells were transfected with Scr or siHIF-2α and empty vector (pcDNA) or Nox2 expression vector (pNox2), followed by analysis of protein expression (*top*) and aconitase activity in cytosolic (*solid*) and mitochondrial (*shaded*) fractions (*bottom)*. (**C**) Intracellular Ca^2+^ concentration (*red*) and calpain activity (*blue*) were determined in cells transfected with Scr or siHIF-2α and either empty vector (pcDNA) or Nox2 vector (pNox2). Ca^2+^ data (mean ± SEM) were obtained from 122 cells. (**D**) HIF-α isoform expression (*top*) and calpain activity (*bottom)* were analyzed in cells transfected with Scr or transfected with siHIF-2α and left untreated (Con) or treated with either 3 µM ionomycin (Ion) or 1.5 µM thapsigargin (Thp). Data are presented as mean ± SEM, *n* = 5-6 independent experiments; \*\**p* < 0.01. (**E**) The mechanism by which HIF-1α negatively regulates HIF-2α and Sod2 expression is shown.

Prior studies revealed that in PC12 cells exposed to intermittent hypoxia, oxidative stress leads to increased intracellular Ca^2+^ levels that trigger HIF-2α degradation by calpains, which are Ca^2+^-dependent proteases (18). Since HIF-1α-deficiency resulted in a reduced intracellular redox state (Figure 6B), we hypothesized that this would result in decreased calpain activity and increased HIF-2α expression. To test this possibility, intracellular Ca^2+^ concentration was analyzed in PC12 cells co-transfected with an expression vector encoding yellow fluorescent protein (YFP) and siHIF-1α. YFP allowed identification of cells that were transfected with siHIF-1α for analysis. Cells transfected with siHIF-1α, but not with scrambled siRNA, manifested decreased intracellular Ca^2+^ levels and decreased calpain activity, and these effects were reversed by forced expression of Nox2 (Figure 6*C*), which normalized the redox state (Figure 6B). Treating siHIF-1α- treated cells with ionomycin or thapsigargin to increase intracellular Ca^2+^ led to increased calpain activity and restored HIF-2α to control levels (Figure 6D). By contrast, PC12 cells transfected with a HIF-1α expression vector exhibited increased intracellular Ca^2+^ and calpain activity, and decreased HIF-2α levels whereas ALLN, a calpain inhibitor, restored HIF-2α protein levels (Figure S6). Thus, loss of Nox2 expression in HIF-1α-deficient cells impairs oxidant- and Ca^2+^-dependent HIF-2α degradation (Figure 6E).

## Discussion

Our results have delineated a previously uncharacterized functional antagonism between HIF-1α and HIF-2α in CB glomus cells and AM chromaffin cells, which regulates hypoxic sensing and is essential for cardiovascular and respiratory homeostasis. CB responses to hypoxia were impaired in *Hif1*α^+/-^ mice and were enhanced in *Hif2*α^+/-^ mice, a finding that is consistent with prior studies (15, 16). We further demonstrate that partial deficiency of either HIF-1α or HIF-2α also alters hypoxic sensing by AM chromaffin cells. Altered hypoxic sensing by the CB and AM was associated with impaired cardio-respiratory homeostasis that was manifested by loss of ventilatory acclimatization to chronic hypoxia in *Hif1*α^+/-^ mice, and by increased BP and respiratory arrhythmias in *Hif2*α^+/-^ mice. Additional studies are warranted to investigate whether HIF-1α or HIF-2α loss-of-function alters the responses of other paraganglionic cells to hypoxia (26).

A major finding of the present study is that the systemic effects of partial loss of either HIF-1α or HIF-2α were associated with increased expression of the other isoform in CB and AM. HIF-1α inhibition in *Hif2α*^+/−^ mice by digoxin treatment, or HIF-2α inhibition in *Hif1α*^+/−^ mice by 2-methoxyestradiol treatment, restored normal CB and AM responses to hypoxia and cardio-respiratory homeostasis. In WT mice, selective HIF-1α inhibition by digoxin treatment led to increased HIF-2α levels, whereas selective HIF-2α inhibition by 2-methoxyestradiol treatment led to increased HIF-1α protein levels, with profound effects on CB and AM function, recapitulating the phenotype seen in *Hif1α*^+/−^ and *Hif2α*^+/−^ mice, respectively. DH mice, which were equally deficient in HIF-1α and HIF-2α, exhibited hypoxic responses and cardio-respiratory function that were similar to WT littermates. Our results demonstrate the existence of functional antagonism between HIF-1α and HIF-2α, and indicate that a fine balance between HIF-α subunits, rather than their absolute levels, is critical for appropriate O_2_ sensing by the CB and AM and for cardio-respiratory homeostasis.

Although HIF-1 and HIF-2 regulate many of the same genes in many tissues (27), recent studies revealed that in the CB and AM they activate expression of gene products with opposing functions that regulate the redox state (11, 17, 18). In the present study, we have demonstrated that imbalanced expression of HIF-1α and HIF-2α alters the intracellular redox state in the CB and AM. *Hif2α*^+/−^ mice exhibited an oxidized, whereas *Hif1α^+/-^* mice displayed a reduced, intracellular redox state. HIF-1α–dependent expression of the pro-oxidant enzyme Nox2 contributed to an oxidized redox state in the CB and AM of *Hif-2α*^+/−^ mice, whereas HIF-2α–dependent expression of the antioxidant enzyme Sod2 contributed to the reduced redox state in the CB and AM of *Hif1α^+/-^* mice. Normalizing the redox state alone was sufficient to normalize the CB and AM responses to hypoxia in *Hif1α^+/-^* and in *Hif2α^+/-^* mice. Thus, complex feed-forward and feed-back relationships between HIF-α isoforms and redox regulatory enzymes establish the redox state, which in turn determines the set point for O_2_ sensing in the CB and AM (Figure 7). Signaling mechanisms including K^+^ channels, Ca^2+^ signaling, mitochondrial metabolism, biogenic amines and gas messengers have been implicated in responses of the CB and AM to hypoxia (19, 28). Further studies are required to determine whether HIF-dependent redox regulation impacts these signaling mechanisms.

**Figure 7.**
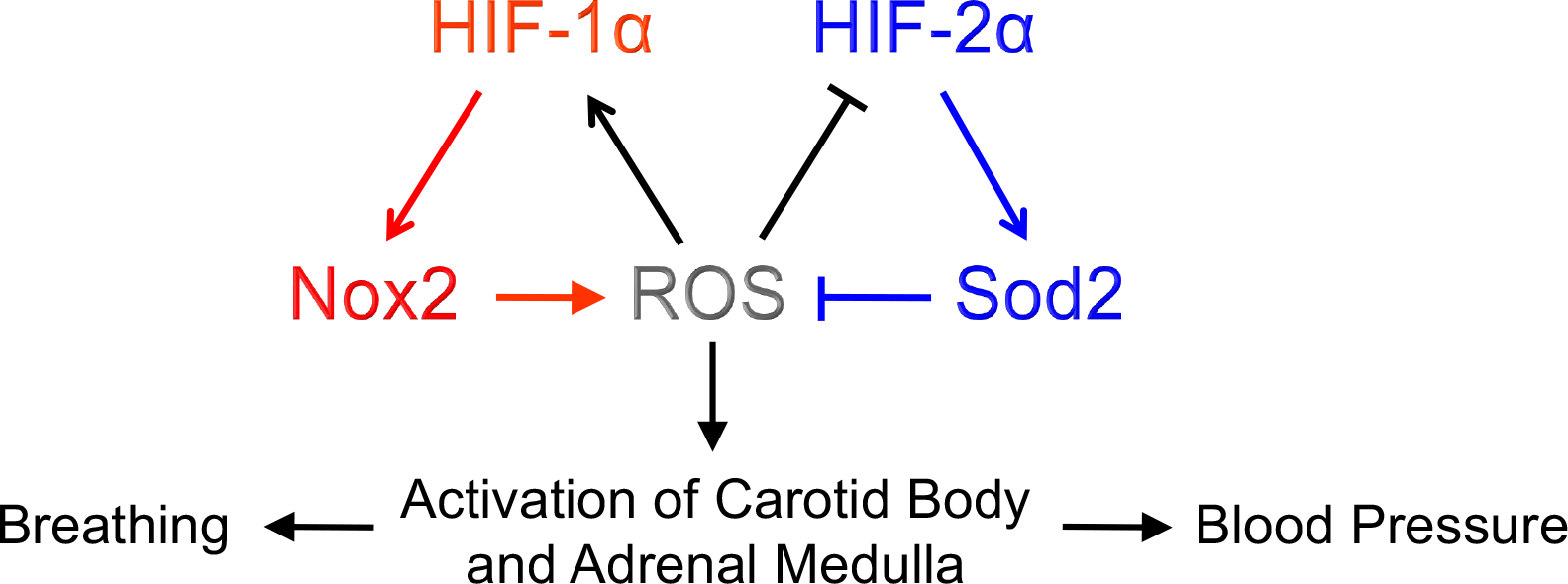
Mutual antagonism between HIF-1α and HIF-2α controls redox status in CB and AM and regulates breathing and BP. The reciprocal positive relationship between HIF-1α and ROS and the reciprocal negative relationship between HIF-2α and ROS are shown.

Oxidant-dependent activation of mTOR signaling and inhibition of prolyl hydroxylase activity, which have been shown to increase HIF-1α protein synthesis (24) and stability (25), respectively, mediated the increased HIF-1α expression in HIF-2α- deficient PC12 cells. On the other hand, loss of oxidant-dependent Ca^2+^-activated calpain activity was responsible for the increased HIF-2α levels in HIF-1α-deficient PC12 cells. Similar mechanisms are likely operative in CB and AM of *Hif1α^+/-^* and *Hif2α^+/-^* mice. Additional studies are required to investigate the mechanism by which HIF-1α deficiency leads to decreased intracellular Ca^2+^ levels and to identify the mechanisms by which calpains recognize and degrade HIF-2α.

Our data indicate that mutual antagonism between HIF-1α and HIF-2α establishes the set point for O_2_ sensing in the CB and AM, which in turn is important for maintaining cardio-respiratory homeostasis. Emerging evidence suggests that hypersensitivity of the CB and AM to hypoxia has an adverse impact on physiological systems. For instance, patients with obstructive sleep apnea exhibit an augment CB response to hypoxia, increased NE levels and BP, and breathing abnormalities (29). Simulating recurrent apnea by exposing rodents to intermittent hypoxia produces an imbalance in HIF-α isoforms manifested by increased HIF-1α (30) and decreased HIF-2α (18) expression in the CB and AM. Intermittent hypoxia-exposed rodents exhibit oxidative stress, increased Nox2 and decreased Sod2 levels, augmented hypoxic responses by the CB and AM, hypertension, and breathing abnormalities (30–32), which recapitulate the phenotype of *Hif-2α*^+/-^ mice. Remarkably, either blockade of changes in HIF-α isoforms or normalization of the intracellular redox state by antioxidants reverses the effects of intermittent hypoxia in rodents (18, 29, 30–33). Thus, correcting the changes in intracellular redox state by modulation of either HIF-1α or HIF-2α levels may represent a novel therapeutic approach for the treatment of cardio-respiratory abnormalities caused by obstructive sleep apnea. Further studies are needed to explore whether altered HIF-α isoform expression and redox state in the CB, AM, or other neuroendocrine cells are involved in other cardio-respiratory diseases. Finally, Belzutifan, a HIF-2α-selective inhibitor has been approved by the U.S. FDA for the treatment of cancer in patients with the von Hippel-Lindau syndrome (34). The results of the present study suggest the possibility that this therapeutic strategy might lead to an imbalance between HIF-1α and HIF-2α in CB and AM, which could result in increased BP and respiratory abnormalities.

## Material and Methods

### Mice

Animal experiments were approved by the University of Chicago Institutional Animal Care and Use Committee and performed by individuals blinded to genotype using male, age-matched *Hif1α^+/-^* and *Hif2α^+/-^* mice and their WT littermates (10, 11).

### Measurements of breathing and BP

Ventilation and O_2_ consumption (*VO2*) were monitored by whole body plethysmography and BP was measured by tail cuff in non-anesthetized mice (16, 30). *Hif2a*^+/-^ mice were anesthetized (70 mg/kg ketamine and 7 mg/kg xylazine, I.P.), the carotid sinus nerves were transected bilaterally, and mice were allowed to recover from surgery for 1 week. Ventilation and BP were analyzed before and after sinus nerve transection. Tidal volume (*V_T_*) and minute ventilation (*V_E_*) were normalized to body weight. Breathing variability was analyzed by Poincarè plots. Breath-to-breath intervals were analyzed as described (16).

### Exposure to hypobaric hypoxia

Non-anesthetized mice were exposed to hypobaric hypoxia as described (15). Following baseline measurements of breathing, mice were placed in a hypobaric chamber (0.4 atmospheres) for 72 hours, and then respiratory responses to acute hypoxia were determined in non-anesthetized mice by whole body plethysmography. Mice were then anesthetized, CBs were harvested and CB responses to hypoxia were determined.

### CB neural activity

Activity from the CB was recorded *ex vivo* as previously described (16, 30). Action potentials (2-3 active units) were recorded from a sinus nerve bundle using a suction electrode. Units were selected based on the height and duration of individual action potentials using a spike discrimination program (Spike Histogram Program, Power Laboratory, AD Instruments). The *P*2 and *P_CO_* of the superfusion medium were measured by a blood gas analyzer (ABL 5, Radiometer). CB activity was averaged during 3 minutes of baseline and 3 minutes of gas challenge and expressed as impulses per second.

### Measurement of plasma NE

Blood was collected from anesthetized mice by cardiac puncture and plasma NE was quantified by HPLC with electrochemical detection using dihydroxybenzylamine as an internal standard (16, 30).

### Measurements of catecholamine secretion

AM chromaffin cells were plated on collagen-coated cover slips in F-12K medium with 1% fetal bovine serum, insulin-transferrin-selenium, and 1% penicillin/streptomycin/glutamine under 7% CO_2_ and 20% O_2_ for 24 hours at 37°C. Catecholamine secretion from individual cells was monitored by carbon fiber amperometry (21). The number of secretory events and the amount of catecholamine secreted per event were analyzed and the data were expressed as total catecholamine molecules secreted.

### Fura-2 imaging

Intracellular Ca ^2+^ levels were quantified in PC12 cells using Fura-2-AM. Transfected cells were identified by yellow fluorescence. Image intensity at 340 nm was divided by 380-nm image intensity to obtain the ratiometric image. Ratios were converted to free intracellular Ca^2+^ using calibration curves constructed *in vitro* by adding Fura-2 (50 μM, free acid) to solutions containing known concentrations of Ca^2+^.

### Reverse-transcription quantitative real-time PCR

HIF-1α, HIF-2α, Nox2, and Sod2 mRNA levels were analyzed by quantitative real-time RT-PCR using SYBR Green (18) normalized to the 18*S* rRNA signal. The following primers were used: HIF-1α: CCACAGGACAGTACAGGATG, TCAAGTCGTGCTGAATAATACC; HIF-2α: TACGAAGTGGTCTGTGGGCAATCA, TCAGCTTGTTGGACAGGGCTATCA; Nox2 (rat): GTGGAGTGGTGTGTGAATGC, TTTGGTGGAGGATGTGATGA; Nox2 (mouse): AGCTATGAGGTGGTGATGTTAGTGG, CACAATATTTGTACCAGACA- GACTTGAG; Sod2: GGCCAAGGGAGATGTTACAGC, GGCCTGTGGTT- CCTTGCAG; and 18S rRNA: CGCCGCTAGAGGTGAAATTC, CGAACCTCCG- ACTTTCGTTCT.

### Cell culture

PC12 cells were cultured in Dulbecco’s Modified Eagle’s medium supplemented with 10% horse serum, 5% fetal bovine serum, penicillin (100 U/ml), and streptomycin (100 μg/ml) under 90% air and 10% CO_2_ at 37°C. Cells were placed in antibiotic free medium and serum starved for 16 hours prior to experiments in order to avoid confounding effects of serum on the expression of HIF-1α or HIF-2α. Cells were pre-treated for 30 minutes with drug or vehicle. PC12 cells were transfected with DNA using Lipofectamine (Gibco). HIF-1α, HIF-2α, Nox2, Sod2 and scrambled siRNAs were purchased from Santa Cruz.

### IB assays

Analysis of HIF-1α, HIF-2α, HIF-1β, Nox2 and Sod2 protein in cell or tissue lysates was performed as described previously (17, 18, 24). The following primary antibodies were used: HIF-1α, HIF-2α, HIF-1β, and prolyl-hydroxylated-HIF-1α (Novus Biologicals); Nox2 (Santa Cruz); phosphorylated and total mTOR (Cell Signaling); Sod2 (Millipore); and α-tubulin (Sigma).

### Enzyme assays

Aconitase, Nox2, and calpain activities were determined as described (17, 18). Sod2 activity was measured using an assay kit (Dojondo Molecular Technologies). Protein levels were determined by Bradford assay kit (Bio-Rad).

### Chemicals

Manganese (III) tetrakis (1-methyl-4-pyridyl) porphyrin pentachloride (MnTMPyP) and digoxin were purchased from Alexis Biochemicals and Sigma-Aldrich, respectively.

### Data analysis and statistics

Data were analyzed with either one-way ANOVA or two-way ANOVA with repeated measures followed by Tukey’s test. Results are reported as mean ± SEM. *P* values of < 0.05 were considered significant. Unless otherwise stated, *n* refers to independent experiments and individual biochemical assays were performed in triplicate.

## Acknowledgments

We thank Kathy Griendling of Emory University for Nox2 vector and Frederick E. Domann of the University of Iowa for Sod2 vector. This research was supported by NIH grants HL76537, HL90554, and HL86493 to N.R.P. and by funds from the Johns Hopkins Institute for Cell Engineering to G.L.S., who is the C. Michael Armstrong Professor at the Johns Hopkins University School of Medicine.

---

**Figure S1.**
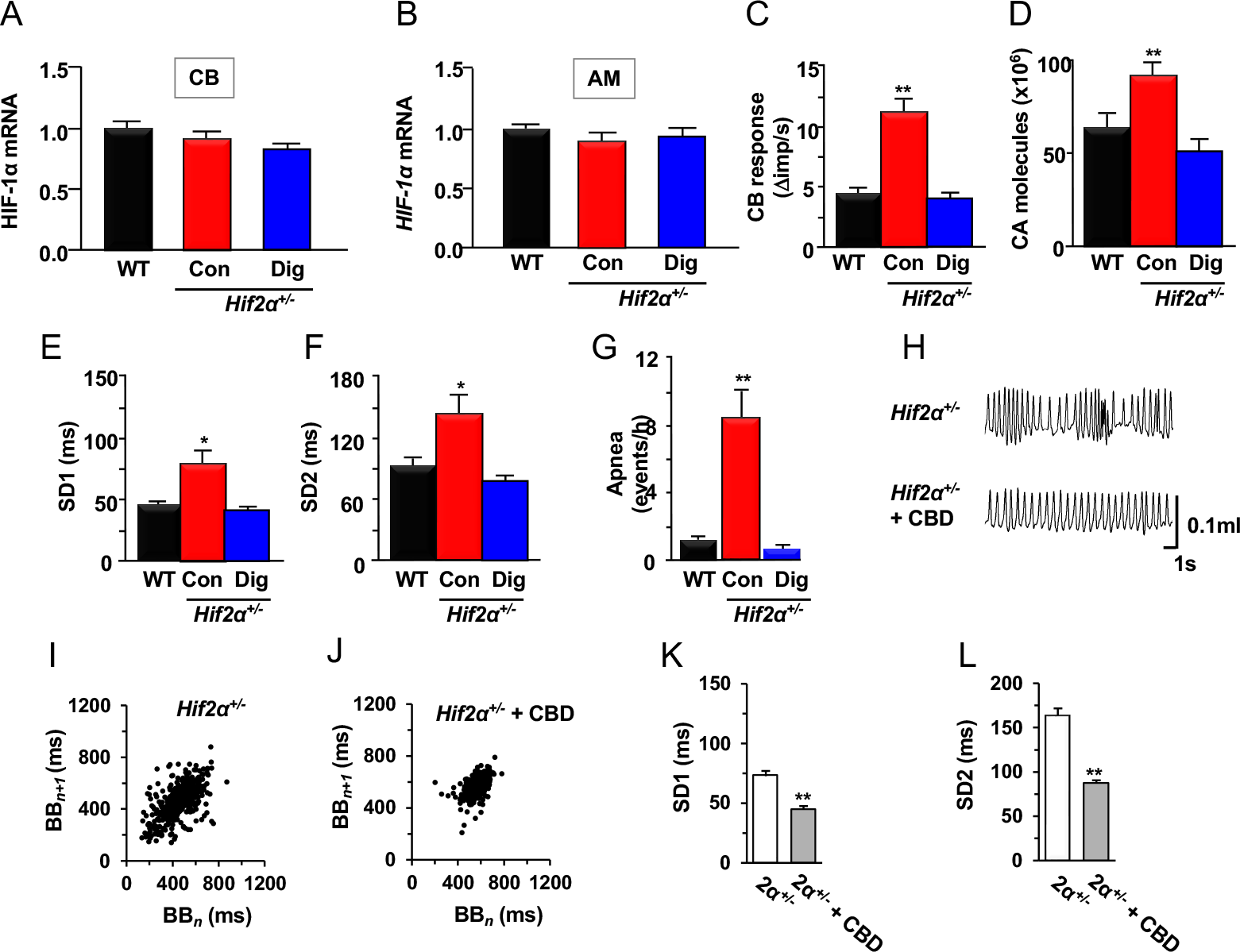
Analysis of *Hif2α^+/−^* mice. The following groups were studied: wild type mice (WT), *Hif2α^+/−^* untreated control mice (Con), and *Hif2α^+/−^* mice treated with digoxin (Dig). (**A** and **B**) HIF-1α mRNA levels in the carotid body (CB) and adrenal medulla (AM) were determined by RT-qPCR. (**C**) CB response to hypoxia (hypoxia – baseline = change in impulses/sec [Δ imp/s]); *n* = 12-15 fibers from each group. (**D**) Catecholamine (CA) secretion from AM chromaffin cells presented as total CA molecules secreted per cell; *n* = 18-22 cells in each group. (**E** and **F**) Quantitative analysis of breathing was performed. Mean standard deviation 1 (SD1; data points from the ascending 45° line; **E**) and mean SD2 (data points from the line orthogonal to SD1; **F**) derived from Poincarè plot analysis of breath-to-breath intervals shown in Figure 1H. (**G**) Apnea index for data derived from Figure 1I. (**H**) Breathing pattern in *Hif-2α*^+/−^ mice with intact CB and after CB denervation (CBD). (**I-L**) Quantitative analysis of breathing was performed using Poincarè plots (**I-J**) to derive standard deviations (**K-L**). Data are presented as mean ± SEM, *n* = 7-8 mice per group; \**p* < 0.05, \*\**p* < 0.01.

**Figure S2.**
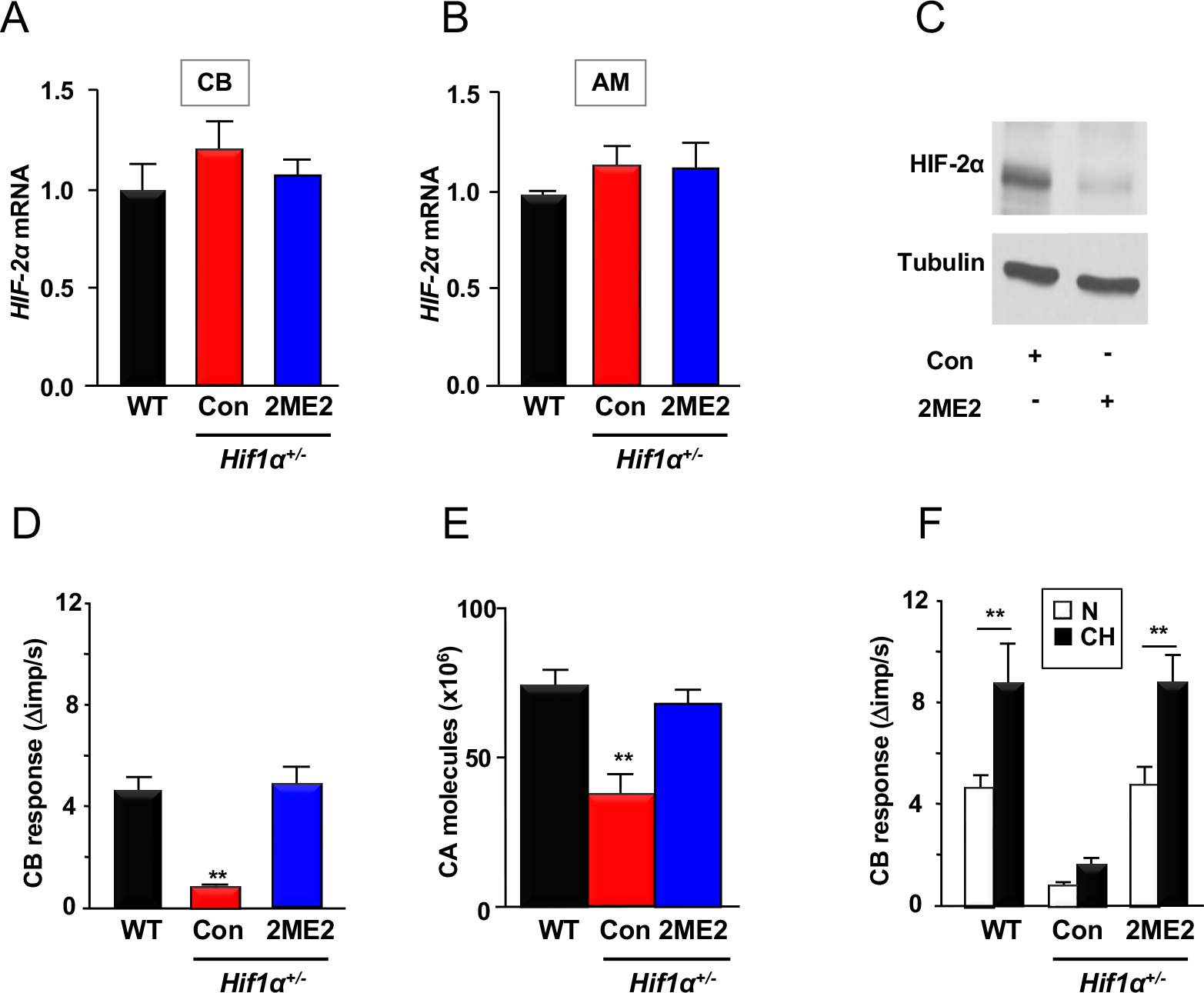
Analysis of *Hif1α^+/−^* mice. The following groups were studied: WT mice, *Hif1α^+/−^* untreated control mice (Con), and *Hif1α^+/−^* mice treated with 2-methoxyestradiol (2ME2). (**A** and **B**) HIF-2α mRNA levels in CB (**A**) and AM (**B**) were determined by RT-qPCR. (**C**) HIF-2α protein expression in PC12 cells treated with 2ME2 (10 μM) as compared to vehicle-treated control cells (Con). (**D**) CB response to hypoxia (*n* = 11-16 fibers per group). (**E**) CA secretion from AM chromaffin cells (*n* = 18-22 cells in each group). (**F**) CB responses to hypoxia in mice exposed to normoxia (N) or chronic hypoxia (CH); *n* = 15-18 fibers per group. Data are presented as mean ± SEM, *n* = 6-8 mice per group; \*\**p* < 0.01.

**Figure S3.**
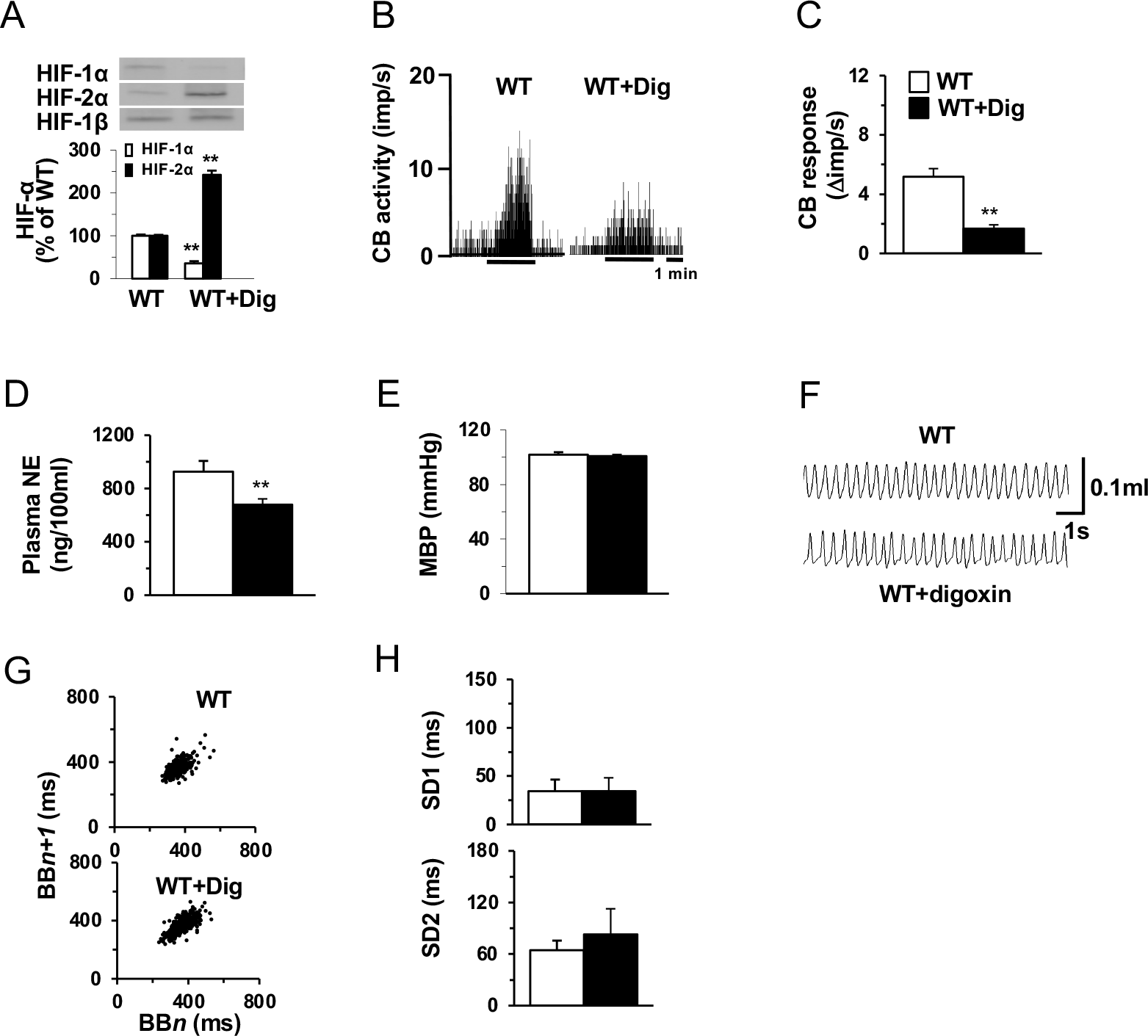
Effect of digoxin in wild type (WT) mice. (**A**) Analysis of HIF-1α, HIF-2α and HIF-1β levels in AM of digoxin-treated WT mice (WT+Dig) relative to untreated WT mice are shown. (**B**) CB responses to hypoxia (*P_O_*_2_ = 40 mm Hg for 3 minutes; black bar under tracing) are shown. Integrated sensory activity (CB activity) is presented as impulses per second (imp/s). (**C**) CB responses to hypoxia (*n* = 11-14 fibers per group). (**D** and **E**) Plasma norepinephrine levels (NE; **D**) and mean blood pressure (MBP; **E**) are shown. (**F-H**) Analysis of breathing, including breathing pattern (**F**), Poincarè plots of breath-to-breath interval (**G**), and derived standard deviations (**H**). Data are presented as mean ± SEM, *n =* 6 mice per group; \*\**p* < 0.01.

**Figure S4.**
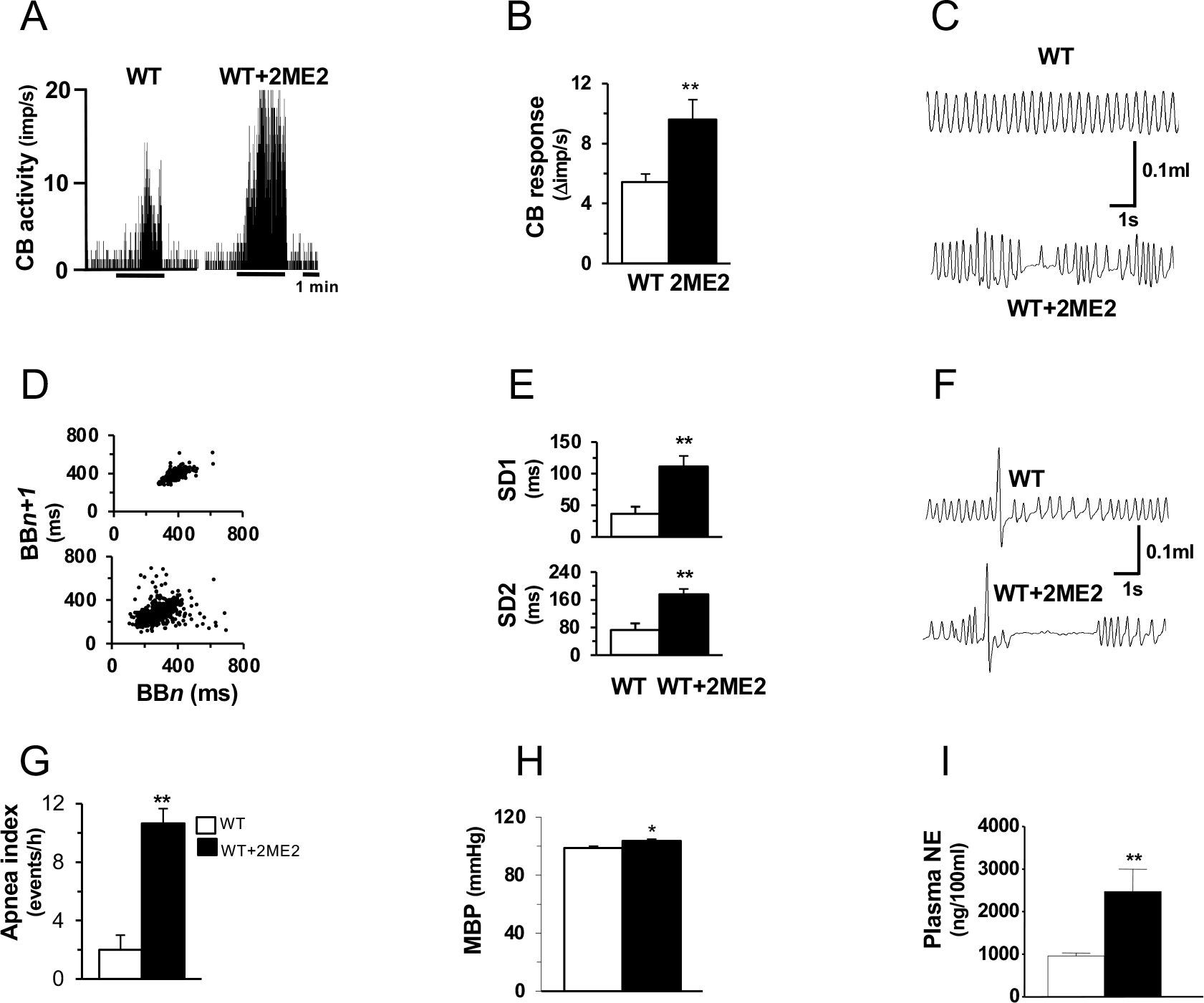
Effect of 2-methoxyestradiol treatment of WT mice. (**A** and **B**) Analysis of CB responses to hypoxia, including single-fiber neural activity recording (**A**) and mean data (**B***; n* = 14-17 fibers per group). (**C-G**) Analysis of breathing, including breathing pattern (**C**), Poincarè plots (**D**) and derived standard deviations (**E**), representative post-sigh apneas (**F**), and apnea index (**G**). (**H** and **I**) Analysis of MBP (**H**) and NE levels (**I**). Data are presented as mean ± SEM, *n* = 6 mice per group; \**p* < 0.05, \*\**p* < 0.01.

**Figure S5.**
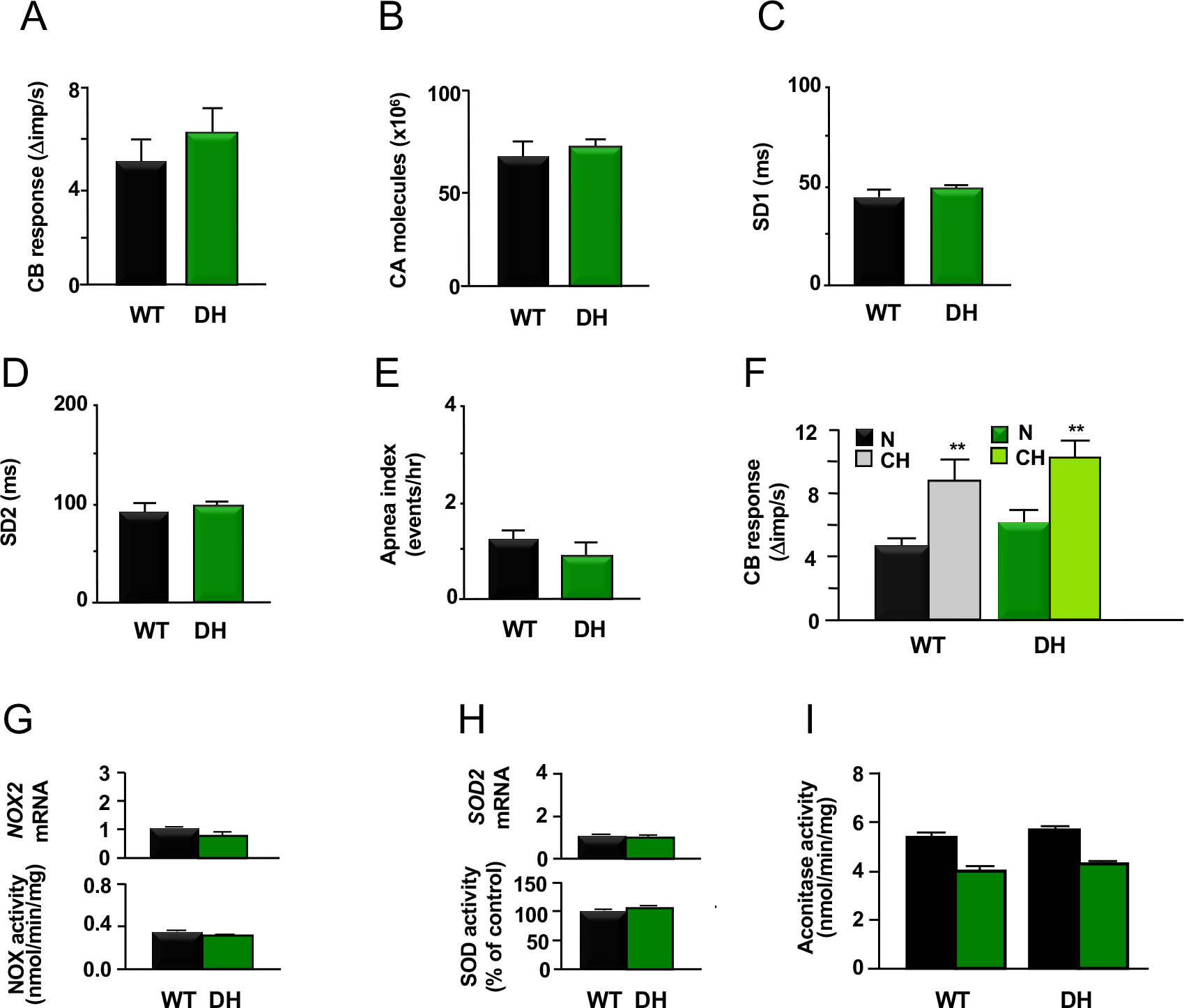
Analysis of DH mice. Two experimental groups were studied: WT mice (*Hif1α^+/+^;Hif2α^+/+^*); and DH mice (*Hif1α^+/-^;Hif2α^+/-^*). (**A**) CB neural activity in response to hypoxia (*n* = 10-14 fibers per group). (**B**) CA secretion from AM chromaffin cells (*n* = 20-24 cells per group). (**C** and **D**) Standard deviations derived from Poincarè plots. (**E**) Apnea index. (**F**) CB neural activity in response to hypoxia in mice exposed to normoxia (N) or chronic hypoxia (CH); *n* = 10-12 fibers per group. (**G-I**) Nox2 and Sod2 mRNA levels and enzyme activities (**G** and **H**), and cytosolic (*black*) and mitochondrial (*green*) aconitase activity (**I**) in AM from WT and DH mice. Data are presented as mean ± SEM; *n* = 6-7 mice per group; \*\**p* < 0.01.

**Figure S6.**
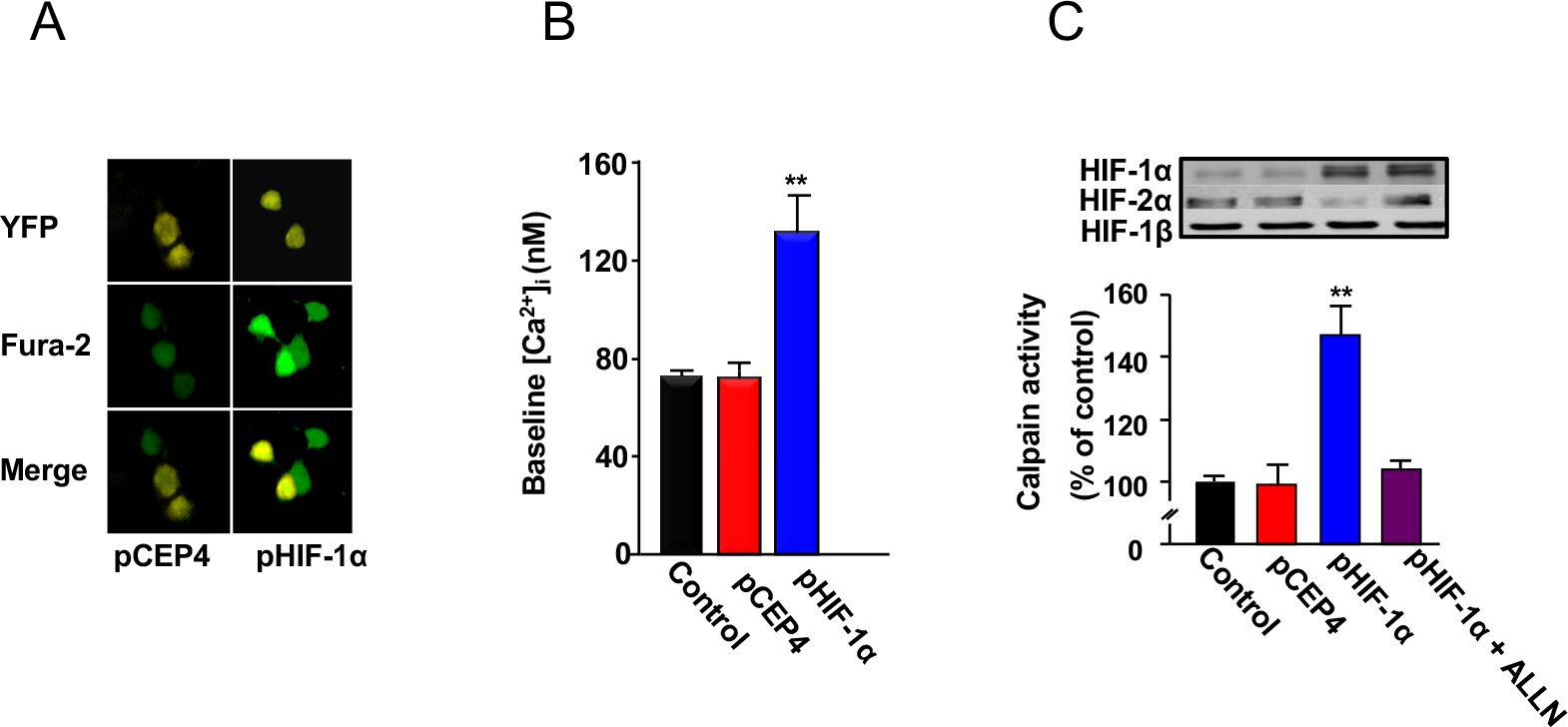
Analysis of changes in intracellular Ca^2+^ levels and calpain activity in response to forced HIF-1α expression in PC12 cells. (**A**) Cells were transfected with a vector encoding yellow fluorescent protein (YFP, *top panels*) and either empty vector (pCEP4) or HIF-1α vector (pHIF-1α). Intracellular Ca^2+^ levels were measured by Fura-2 imaging (*middle panels*) and YFP and Fura-2 images were merged (*bottom panels*). (**B**) Intracellular Ca^2+^ concentration was measured in control (non-transfected) cells or cells transfected with indicated plasmid; *n* = 96-115 cells per group. (**C**) HIF subunit expression (*immunoblots*) and calpain activity (*bar graph*) were determined in control cells and cells transfected with pCEP4 or pHIF-1α, or transfected with pHIF-1α and treated with 10 μM ALLN. Data are presented as mean ± SEM, *n* = 4 independent experiments; \*\**p* < 0.01.

